# Photoperiod-induced neurotransmitter plasticity declines with aging: an epigenetic regulation?

**DOI:** 10.1101/563213

**Authors:** Rory Pritchard, Helene Chen, Ben Romoli, Nicholas C. Spitzer, Davide Dulcis

## Abstract

Neuroplasticity has classically been understood to arise through changes in synaptic strength or synaptic connectivity. A newly discovered form of neuroplasticity, neurotransmitter switching, involves changes in neurotransmitter identity. Chronic exposure to different photoperiods alters the number of dopamine (tyrosine hydroxylase, TH+) and somatostatin (SST+) neurons in the paraventricular nucleus (PaVN) of the hypothalamus of adult rats and results in discrete behavioral changes. Here we investigate whether photoperiod-induced neurotransmitter switching persists during aging and whether epigenetic mechanisms of histone acetylation and DNA methylation may contribute to this neurotransmitter plasticity. We show that this plasticity is robust at 1 and at 3 months but reduced in TH+ neurons at 12 months and completely abolished in both TH+ and SST+ neurons by 18 months. *De novo* methylation and histone 3 acetylation were observed following short-day photoperiod exposure in both TH+ and SST+ neurons at 1 and 3 months while an overall increase in methylation of SST+ neurons paralleled neuroplasticity reduction at 12 and 18 months. Histone acetylation increased in TH+ neurons and decreased in SST+ neurons following short-day exposure at 3 months while the total number of acetylated PaVN neurons remained constant. Reciprocal histone acetylation in TH+ and SST+ neurons suggests the importance of studying epigenetic regulation at the circuit level for identified cell phenotypes. The association of age-dependent reduction in neurotransmitter plasticity and changes in DNA methylation and acetylation patterns in two neuronal phenotypes known to switch transmitter identity suggests mechanistic insights into transmitter plasticity in the aging brain.

**SIGNIFICANCE:** Neurotransmitter switching, like changes in synaptic strength, formation of new synapses and synapse remodeling, declines with age. This age-dependent reduction in transmitter plasticity is associated with changes in levels of DNA methylase and histone deacetylase that imply epigenetic regulation of transcription. A reciprocal pattern of histone acetylation in a single population of neurons that depends on the transmitter expressed emphasizes the value of studying epigenetic mechanisms at the level of cell phenotypes rather than cell genotypes or whole tissue. The findings may be useful for developing approaches for non-invasive treatment of disorders characterized by neurotransmitter dysfunction.

## INTRODUCTION

During embryonic development, genetic programs organize expression patterns of neurotransmitters within specific neuronal networks and activity-dependent mechanisms mediated by calcium signaling enable refinement of neurotransmitter expression (Borodinsky et al., 2004; Rosenberg and Spitzer, 2011). Experience-dependent neurotransmitter plasticity extends beyond this early phase. Induced by natural stimuli during development and in the mature nervous system, it entails addition, loss, or switching of neurotransmitters in a process known as neurotransmitter respecification (Dulcis and Spitzer, 2008; Dulcis et al., 2013, 2017; Spitzer, 2015).

Seasonal changes in photoperiod act as environmental stressors, affecting expression of clock genes that control functions of the circadian pacemaker, the suprachiasmatic nucleus (SCN) of the hypothalamus. SCN neurons make connections to the Paraventricular Nucleus (PaVN) of the hypothalamus that integrates neuroendocrine and autonomic functions (Ferguson et al., 2008). Local PaVN interneurons expressing TH or SST or both make synapses on corticotropin-releasing factor (CRF) cells (Dulcis et al., 2013; Cummings et al., 1983; Kumar, 2007), modulating release of CRF and regulating stress responses (Herman and Rosenmund, 2015). Exposure of adult rats to long-day (19L:5D) or short-day (5L:19D) photoperiods leads to a shift in the number of dopaminergic (DA+TH+) and somatostatin (SST+) neurons in the PaVN, resulting in behavioral changes associated with stress responses (Dulcis et al., 2013). Rats exposed to long-day photoperiods for 1 week exhibit an increase in SST+ neurons and a decrease in TH+ neurons that cause more depressive- and anxiety-like behaviors when tested on the elevated plus maze and forced swim tests, and display a higher titer of stress hormone CRF in their cerebral spinal fluid. Short-day exposure causes the opposite effects on the number of TH+ and SST+ neurons, stress responses, and CRH titer (Dulcis et al., 2013). Presynaptic changes in neurotransmitter identity were matched by changes in post-synaptic receptors in CRF cells, regulating stress responses. Neurotransmitter switching occurs through recruitment of “reserve pool” neurons that adapt circuit function by changing neurotransmitter identity (Dulcis and Spitzer, 2012).

The activity-dependent genetic regulation of photoperiod-induced neurotransmitter plasticity affecting behavior has implications for people who suffer from Seasonal Affective Disorder (SAD) as well as Major Depressive Disorders (Lam et al., 2006, 2016). Due to its seasonal recurrence and equal incidence among men and women, SAD is a serious mental health problem (Kurlansik and Ibay, 2007). Because SAD’s prevalence declines with age, it is of interest to investigate whether photoperiod-induced neurotransmitter plasticity is affected by aging.

We hypothesized that DNA modification has a role in photoperiod-induced plasticity, given that transmitter switching requires chronic neuronal activation (Meng et al., 2018) and that neuronal activity modifies chromatin accessibility and gene expression via epigenetic mechanisms in adult neurons (Su et al., 2017; He et al., 2011). While *de novo* transcriptional regulation has been shown to be the mechanism behind light-induced dopamine/SST transmitter switching (Dulcis et al., 2013), it was unknown whether DNA methylation or histone acetylation are potential environmental links initiating the genetic cascade regulating this form of plasticity. DNA methylation is essential for cellular mechanisms involved in synaptic plasticity, neuronal repair, neuronal survival, and learning and memory (Lardenoije et al., 2015; Fan et al., 2001; Feng et al., 2010; Iskandar et al., 2010). It is generally associated with gene repression (Moore et al., 2013). However, methylation of specific local non-promoter sites can also result in gene activation and hypermethylation of certain genes results in increased expression (Wu et al., 2010; Silva et al., 2008). Histone acetylation makes DNA more accessible to regulatory proteins and transcription factors (Grunstein, 1997). The roles of *de novo* DNA methylation and histone acetylation in neural plasticity and emotional behavior (Oliveira et al., 2012) suggest that these processes could have particular relevance to the mechanism of neurotransmitter respecification in the mature and aging brain.

In this study we provide evidence that neurotransmitter plasticity is age-dependent and discover cell-specific epigenetic patterns of DNA methylation and histone acetylation in response to changes in photoperiod.

## MATERIALS and METHODS

### Animals

Male Long Evans rats (Harlan Industries, Indianapolis, IN) were separated into groups by age (1, 3, 12, and 18 months) and by experimental photoperiod conditions. Animals were exposed for 1 week to controlled light/dark cycles consisting of either a long day [19 hours light: 5 hours dark (19L:5D)], normal day [12 hours light:12 hours dark (12L:12D)] or short day [19 hours light:5 hours dark (19L:5D)]. During photoperiod exposure, rats were singly housed with food and water *ad libitum*. All protocols were in strict adherence to the National Institutes of Health *Guide for the Care and Use of Laboratory Animals* and approved by the IACUC at University of California, San Diego.

### Photoperiod chambers

Wooden photoperiod-chamber boxes were purchased from Actimetrics (Wilmette, IL). Each box was equipped with its own ventilation system, LED lights (134 LUX) set on timers to specific photoperiod conditions and completely sealed off from external room lighting. Animals were allowed to habituate in the chambers for 48 hours on a normal 12L:12D day. Following habituation, animals were exposed to their respective photoperiod conditions for 1 week, as 1 week has been shown to be sufficient time to induce neurotransmitter switching in the adult rat hypothalamus (Dulcis et al., 2013),

### Immunohistochemistry

Immediately after one week of exposure, all animals were perfused during the light phase of their light/dark cycle. Rats were anesthetized with 5% isoflurane (Vet One, Boise, ID), followed by an intraperitoneal (i.p.) injection of sodium pentobarbital (Vortech Pharmaceuticals, Dearborn, MI) at a dose of 150 mg/kg. Animals were then transcardially perfused with 1x PBS until perfusate was clear (250-300 mL), followed by ice-cold 4% paraformaldehyde (MP Biomedicals, Santa Ana, CA) until the neck and hindquarters were rigid (300 mL). Brains were extracted, post-fixed overnight in 4% paraformaldehyde, then cryoprotected in 30% sucrose for 48 hr. Once cryoprotected, brains were flash frozen on dry ice and 30 µm-thick coronal sections were collected in a 1:12 series using a Leica SM2010R microtome. Tissue sections were stored in cryoprotectant (30% glycerol, 30% ethylene-glycol in 1xPBS, Fisher Bioreagents, Fair Lawn, NJ) at −20° C until ready for processing.

Immunohistochemistry for tyrosine hydroxylase (TH) and somatostatin (SST) was performed on free-floating sections using the standard avidin-biotin peroxidase method (Dulcis et al., 2013). Tissue sections were rinsed through a succession of 1x PBS washes, blocked for 30 min at room temperature with 5% normal horse serum (Vector Laboratories, Burlingame, CA) in 0.3% PBS-TX (Triton X-100, Fisher Bioreagents, Fair Lawn, NJ), and incubated in either mouse anti-TH [1:1000] (Millipore, Temecula, CA, lot # 2716631) overnight or goat anti-SST [1:500] (Santa Cruz Biotechnology, Dallas, TX, lot # L0514) for 48 hr at 4°C. Tissue sections were then rinsed through another series of 1x PBS washes and incubated in either biotinylated anti-mouse IgG or anti-goat IgG secondary antibody [1:200] (Vector Laboratories, Burlingame, CA). The signal was amplified using a Vectastain ABC kit (Vector Laboratories, Burlingame, CA) and developed with diaminobenzidine (DAB, Acros Organics, Morris Plains, NJ). Sections were mounted on superfrost slides (Fisher Scientific. Pittsburgh, PA) in 0.2% gelatin (Millipore, Burlington, MA), dried overnight at 21°C, dehydrated and coverslipped with Cytoseal (Thermoscientific, Kalamazoo, MI). Tissue sets stained for TH were counterstained using a Giemsa stain (1:2 solution in 0.1x PBS, Harleco, EMDmillipore, Temecula, CA) to identify the boundaries of the PaVN.

Triple-label immunofluorescence was used to identify levels of methylation and acetylation in TH+ and SST+ neurons of the paraventricular nucleus of the hypothalamus (PaVN). Free-floating tissue sections were rinsed with 1x PBS, blocked with 5% normal horse serum in 0.3% PBS-TX, and labeled with either rabbit anti-DNMT3a [1:500] (Santa Cruz Biotechnology, Dallas, TX) or rabbit anti-AcetylH3 [1:500] (Millipore, Temecula, CA, lot #2153150), along with mouse anti-TH [1:1000] (Millipore, Temecula, CA, lot # 2716631) and goat anti-SST [1:500] (Santa Cruz Biotechnology, Dallas, TX, lot # L0514) for 48 hr at 4°C. Tissue sections were then rinsed through another succession of 1x PBS washes and labeled with species matched secondary antibody AlexaFluor conjugates [1:300]: donkey anti-rabbit 488, donkey anti-goat 555 and donkey anti-mouse 647 (Invitrogen, Carlsbad, CA). Sections were rinsed again in 1x PBS, mounted on superfrost slides in 0.2% gelatin and coverslipped with Fluoromount-G (Southern Biotech, Birmingham, AL).

### Imaging and neuronal quantification

Images of TH+ and SST+ DAB-labeled tissue were captured in bright field with either a Hamamatsu Nanozoomer 2.0HT slide scanner or a Leica DM4 B Stereologer for stereological quantification. Fluorescence microscopy was performed on a Leica TCS DMi8 confocal microscope. System optimized z-stacks were collected and processed using the LAS AF software. Cell counts of epifluorescence immunoreactivity were performed using Adobe Photoshop CC.

### qPCR

Immediately after photoperiod exposure, rat brains were rapidly extracted and the hypothalamus was dissected in an RNAse-free environment and processed for qPCR. Bilateral tissue punches (1.5 mm) were collected from the PaVN and stored in RNAlater (ThermofFisher Scientific, Waltham, MA). Tissue punches were homogenized in Trizol (Thermofisher) using 0.5 mm zirconium oxide beads and a bullet blender (Next Advance Inc., Troy, NY). RNA was purified with Direct-zol kit (Zymo, Irvine, CA) and converted to cDNA using an iScrip kit (Biorad, Hercules, CA) following manufacturer’s instructions. RT-PCR was performed in a Biorad CFX384 using KiCqStart SYBR Green Ready Mix (Millipore-Sigma, St. Louis, MO). RNA expression was normalized to the housekeeping gene GADPH and quantified using the ΔΔCt method. Primer pair (5’-3’) sequences used for each candidate gene are listed below.

*Dnmt1* (F: TTCTCGGCAGGGTATG; R: GTCTACCGACTGGGTGACAGT), *Dnmt3a* (F: ACGCCAAAGAAGTGTCTGCT; R: CTTTGCCCTGCTTTATGGAG), *Dnmt3b* (F:CATAA GTCGAAGGTGCGTCGT; R: ACTTTTGTTCTCGCGTCTCCT), *G9a (F:* GACAACAAG GATGGTGAGGTC; *R*: AGCATGAAGACCCGAACAG), *GCN5* (F: CATCGGTGGGATT TGCTT; R: GTACTCGTCGGCGTAGGTG), *P300* (F: GGGACTAACCAATGGTGGTG; R: ATTGGGAGAAGTCAAGCCTG), *CBP* (F: GACCAAGATGGGGATGACTG; R: CCA CTGATGTTTGCAACTGG), *Housekeeping gene GAPDH* (F: TATGATGACATCAAGG TGG; R: CACCACCCTGTTGCTGTA).

### Experimental Design and Statistical Analysis

We hypothesized that the extent of plasticity would be greater in younger animals and reduced in older animals. Our baseline studies had focused on adult Long Evans male rats, 3 months of age. To extend this work we studied young rats, 1 month of age, and older rats, 12 and 18 months of age. Animals of each age were exposed to short-day, long-day and balanced-day light-dark cycles for one week, in the same photochambers with the same intensity of illumination. Brains were perfused, collected, sectioned and immunostained for neurotransmitter and epigenetic markers. Because 18-month old rats display more than 50-70% mortality before they reach (for rats) this old age, generating a sufficient number of aged rats for our experiments was a limiting step. We solved the problem working more closely with the staff of our approved vendor, Harlan Laboratories, who were able to generate aged rats by setting aside up to 40 rats in order to yield at least 10 aged animals. Normally distributed data were analyzed using two-tailed Student’s t-test and one-way, two-way or repeated measures ANOVA, followed by post-hoc comparison. All data were analyzed with IBM SPSS 24.0 (Chicago, IL) and represented by bar plot with mean and SEM (standard error). Significance in the figures was indicated with * P≤0.05; ** P≤0.01; *** P≤0.001. Exact P and t-test values are reported for each experiment.

## RESULTS

### Photoperiod-induced dopamine plasticity declines with age

We hypothesized that the extent of neurotransmitter plasticity would be greater in younger animals and reduced in older animals. Our previous studies focused on adult male rats, 3 months of age (Dulcis et al., 2013). To extend this work we included the analysis of younger rats, 1 month of age, as well as older rats, 12 and 18 months of age. Animals of each age group were exposed to short-day (5L:19D), long-day (19L:5D), and balanced-day (12L:12D) light-dark cycles for one week. Our findings indicate that there is a decline of dopamine plasticity with aging. To compare changes across photoperiods and age groups we calculated a plasticity index. The mean number of neurons in balanced-day control photoperiod for each age group was normalized to 100% and given a plasticity value of 1; all other mean values corresponding to short- or long-photoperiods were divided by the mean of control photoperiod of their age group. Exposure to short- and long-day photoperiods affected the number of TH+ neurons in the PaVN compared to control animals. Juvenile animals (1-month old) were the most responsive age group, displaying the highest TH plasticity index in response to short-day photoperiod exposure (Figure 1A, 1E), compared to adults (Figure1B, 1E). Long-day exposure significantly reduced the number of TH+ cells in the PaVN of animals at 1 (Figure 1A, 1E) (t_(10)_ = 2.45, p = 0.0087 unpaired), 3 (Figure 1B, 1E) (t_(21)_ = 5.11, p = 0.000023, unpaired) and 12 months of age (Figure 1C, 1E) (t_(12)_ = 4.48, p = 0.0007, unpaired). Conversely, short day exposure significantly increased the number of TH+ cells in the PaVN at 1 (Figure 1A, 1E) (t_(10)_ = 3.25, p = 0.0204, unpaired) and 3 months of age (Figure 1B, 1E) (t_(21)_ = 2.55, p = 0.0204, unpaired). However this effect was lost at 12 and 18 months (Figure 1C, 1D, 1E). Interestingly, rats at 12 months of age showed a significant decrease in dopaminergic neurons only in response to long-day periods but no longer exhibited the increase with short-day exposure (Figure 1C, 1E). In aging rats (18 months), photoperiod induced TH-induced plasticity was completely abolished (Figure 1D, 1E). These findings demonstrate that young animals show greater photoperiod-induced neuroplasticity than aging rats. Moreover, mature rats (3-12 months old) are more susceptible to light-induced dopamine switching in response to a stressful stimulus (long day photoperiod) than they are to a more pleasurable environment (short day photoperiod).

**Figure 1.**
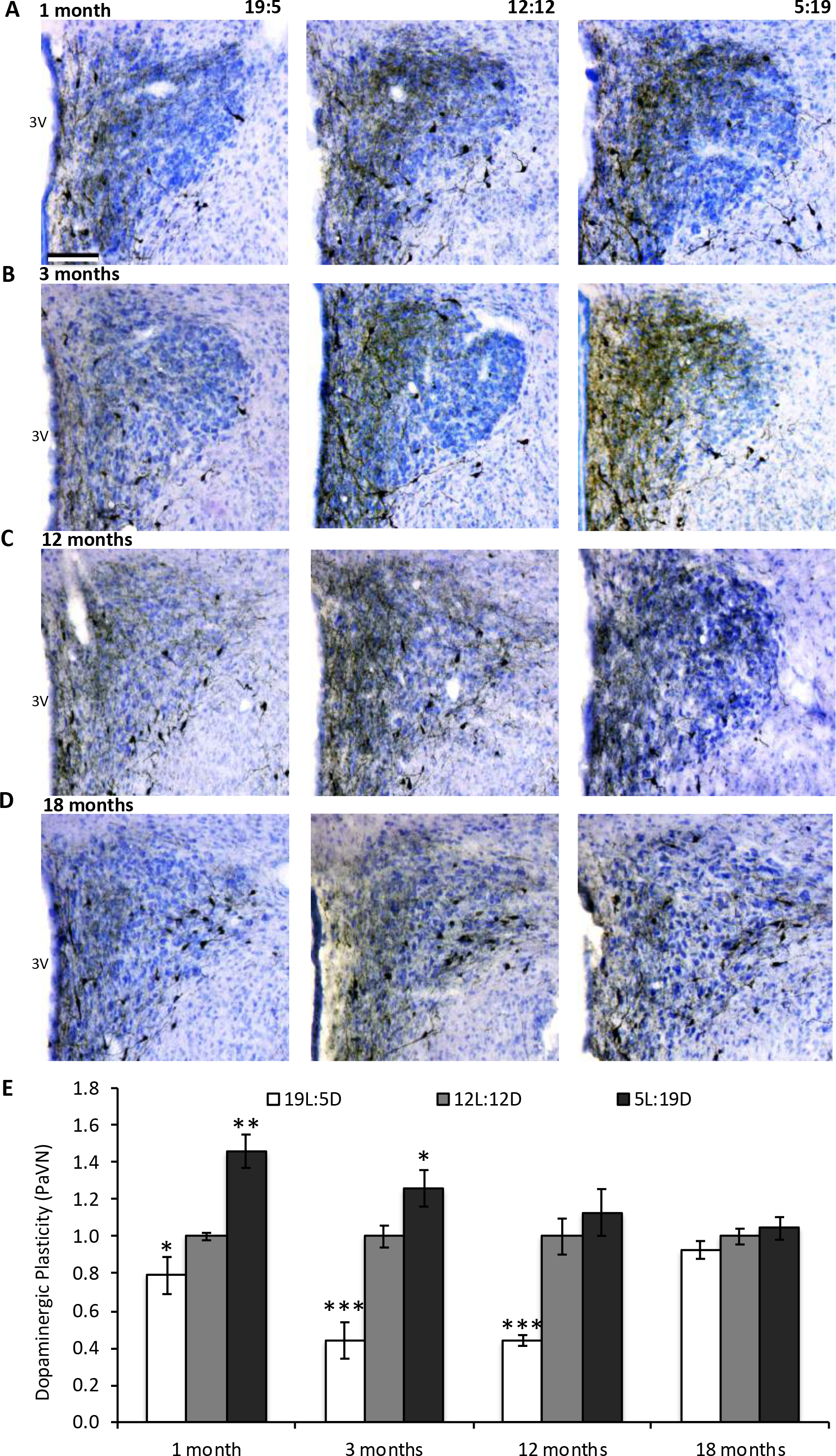
Photoperiod-induced dopamine plasticity declines with age. (**A-D**) Bright-field images of PaVN sections immunostained with tyrosine hydroxylase (TH) antibody labeled with DAB and counterstained with Giemsa. Exposure to short-day (5L:19D, *right*) and long-day (19L:5D, *left*) photoperiods for 1 week alters the number of TH+ neurons in the PaVN relative to control animals (12L:12D, *middle*) in an age-dependent manner. (**E**) The number of TH+ neurons in the PaVN was compared among age groups using a normalization index of dopaminergic plasticity. Younger animals at 1 (N = 6) and 3 months (N = 10) showed a higher responsiveness to photoperiod, exhibiting a marked increase in number of TH+ neurons when exposed to a short-day light cycle and a significant decrease in number when exposed to a long-day cycle. At 12 months of age (N = 8), rats displayed a significant decrease in TH+ neurons following long-day light cycle relative to controls, but no change when exposed to a short-day light cycle. At 18 months (N = 6), the photoperiod effect was lost and there were no changes in the number of TH+ neurons across photoperiods. Scale bar, 100 µm. Values are mean ± SEM. **p= ≤ 0.01). 3V, third ventricle.

### Photoperiod-dependent SST plasticity is also abolished with aging

Consistent with previous studies (Dulcis et al., 2013), short- and long-day exposure significantly influenced the number of SST+ neurons in the PaVN in an inverse relationship with the number of TH+ neurons. Long-day exposure significantly increased the number of SST+ cells in the PaVN in animals at 1 (t_(10)_ = 5.31, p = 0.0003, unpaired), 3 (t_(12)_ = 3.55, p = 0.0039, unpaired) and 12 (t_(12)_ = 2.84, p = 0.0146, unpaired) months of age (Figure 2A, 2B, 2C, 2E). Short-day exposure had the opposite effect in animals of these three age groups (1 month, t_(10)_ = 2.99, p = 0.010, unpaired; 3 months, t_(10)_ = 3.52, p = 0.0055, unpaired; 12 months, t_(12)_ = 5.78, p = 0.000043, unpaired), in which the number of SST+ cells in the PaVN was greatly reduced as the number of TH+ neurons was increased. Unlike photoperiod-dependent TH plasticity, short-day exposure in 12-month old animals induced SST neuroplasticity (Figure 2C, 2E). As observed for dopamine neurons, photoperiod-induced SST switching was completely abolished in animals at 18 months of age (Figure 2D, 2E). These data indicate that photoperiod-induced transmitter plasticity is age-dependent.

**Figure 2.**
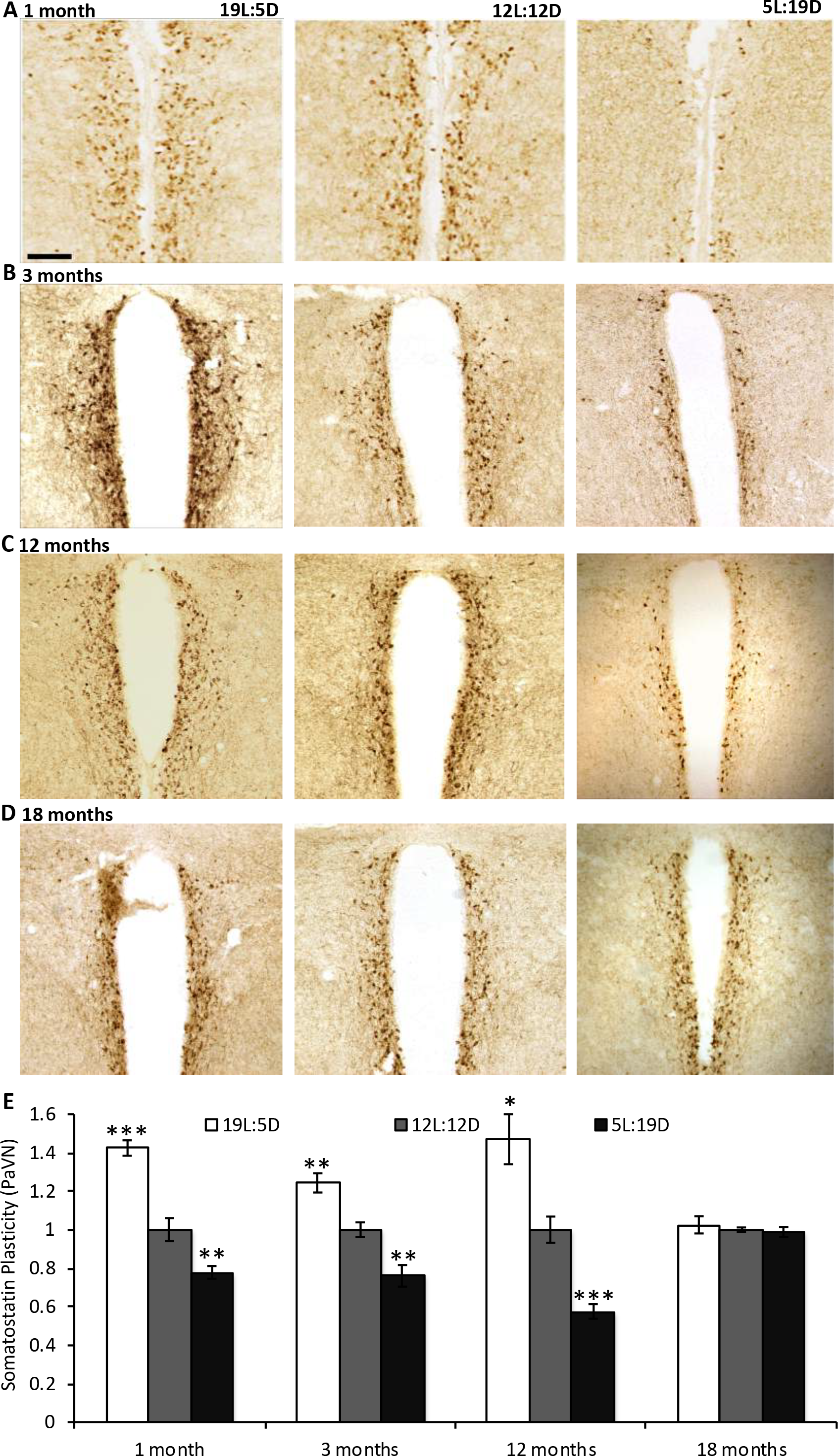
Photoperiod-induced somatostatin plasticity declines with age. (**A-D**) Bright-field images of PaVN sections immunostained with somatostatin (SST) antibody labeled with DAB. Exposure to short-day (5L:19D, *right*) and long-day (19L:5D, *left*) photoperiods for 1 week alters the number of SST+ neurons in the PaVN relative to control animals (12L:12D, *middle*) in an age-dependent manner. (**E**) The number of SST+ neurons in the PaVN was compared among age groups using a normalization index of somatostatin plasticity. Animals at 1 (N = 6), 3 (N = 8) and 12 (N = 8) months of age showed photoperiod-dependent neurotransmitter switching in an inverse relation with respect to TH plasticity. As seen for TH expression, the effect was lost at 18 months of age (N = 6) showing no change in the number of SST+ neurons across all photoperiods. Scale bar, 100 µm. Values are mean ± SEM. **p= ≤ 0.01); ***p= ≤ 0.001).

### Patterns of Dnmt3a and Acetyl H3 mRNA change with photoperiod exposure

To investigate the epigenetic mechanisms that may contribute to photoperiod-induced neurotransmitter respecification, we performed qRT-PCR of hypothalamic paraventricular tissue to identify changes in transcriptional regulation of methyl- and acetyl-transferases, *Dnmt1*, *Dnmt3a*, *Dnmt3b*, *G9a*, *GCN5*, *P300, CBP*, which have been shown to be regulated in association with various forms of neuroplasticity (Laplant et al., 2010; Maze et al., 2010; Wu et al., 2017; Park et al., 2012). We measured strong mRNA upregulation of all methyltransferases in response to altered photoperiods (Fig. 3). Acetyltransferases displayed regulation in response to altered photoperiods that in many cases was not significant (Fig.3).

**Figure 3.**
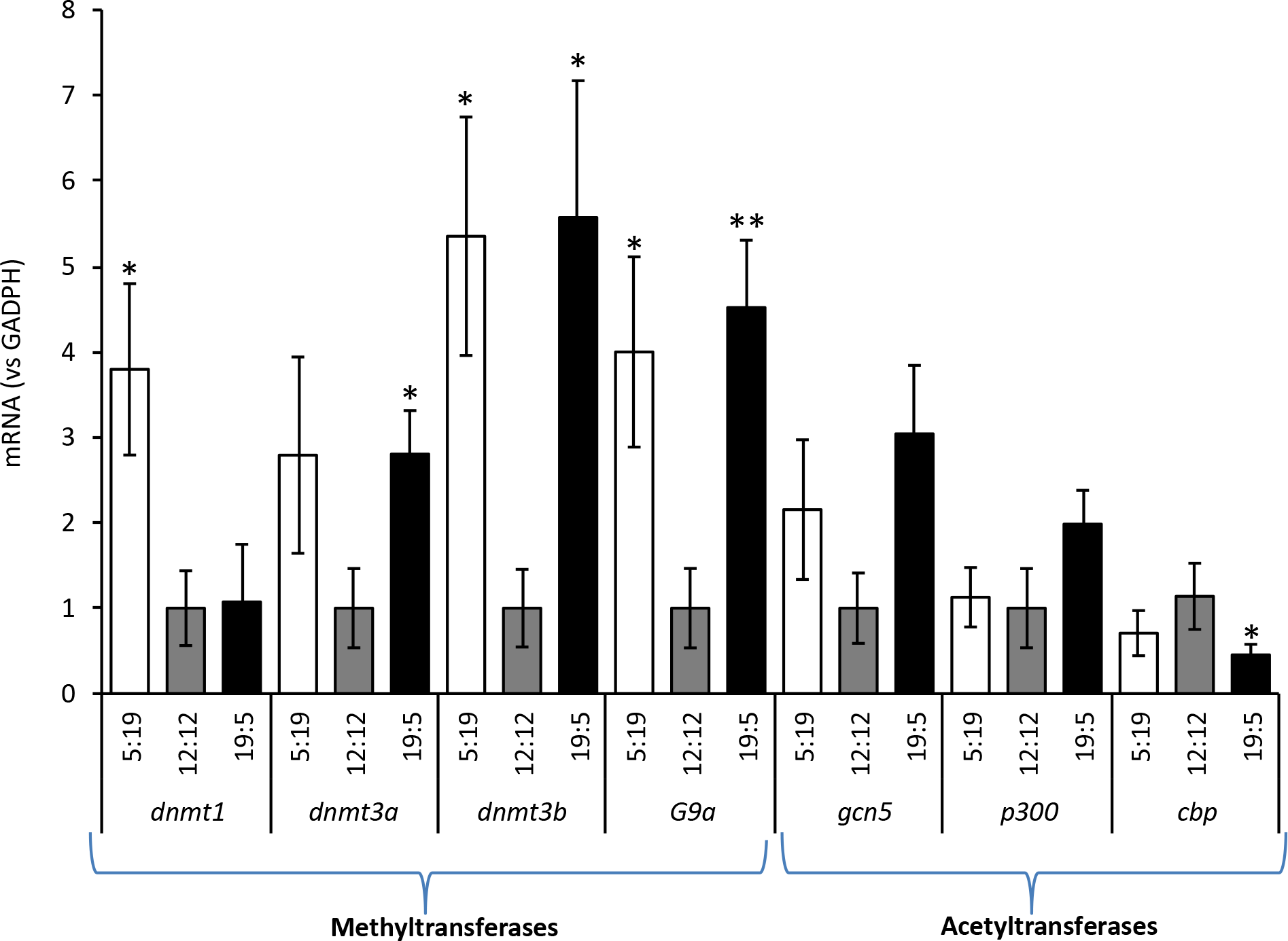
Transcriptional regulation of transferases in response to altered photoperiod. Photoperiod regulates the mRNA levels of specific DNA methyl- and histone acetyl-transferases in the PaVN. Quantification shows significant (vs GADPH) changes of DNA transferase transcripts compared to balanced (12L:12D) photoperiod (N=5 rats per group). Values are mean ± SEM. *Dnmt1 (5:19, t_(8)_* unpaired = 2.55, p = 0.034); *Dnmt3a (19:5, t_(8)_* unpaired = 2.61, p = 0.031); *Dnmt3b (5:19, t_(8)_* unpaired = 2.97, p = 0.018*; 19:5, t_(8)_* unpaired = 2.75, p = 0.025); *G9a (5:19, t_(8)_* unpaired = 2.48, p = 0.038*; 19:5, t_(8)_* unpaired = 3.84, p = 0.005); *CBP (19:5, t_(8)_* unpaired = 2.77, p = 0.024).

### DNA methylation becomes photoperiod-independent with aging

We then investigated protein expression of a candidate methyl-transferase, DNMT3a, in TH+ and SST+ neurons that display neurotransmitter plasticity in the PaVN and compared the results across age groups and photoperiods (Figure 4A). Animals at 1 (TH, t_(4)_ = 2.87, p = 0.0452, unpaired; SST, t_(4)_ = 1.00, p = 0.3739, unpaired) and 3 months (TH, t_(4)_ = 2.93, p = 0.0424, unpaired; SST, t_(4)_ = 2.90, p = 0.0439, unpaired) had a significant increase in the number of both TH+ and SST+ cells that colocalized with DNTM3a in the PaVN following short-day exposure only (Figure 4B, 4C). Long-day exposure showed no difference in DNA methylated TH+ or SST+ neurons from balanced-day controls. The photoperiod effect in methylating DNA of TH+ and SST+ cells in the PaVN was lost in rats at 12 and 18 months of age and was replaced by a marked overall increase in methylation of SST+ cells that was photoperiod-independent (Figure 4D, 4E). The reciprocal aging effect on the Dnmt3a-methylation level of SST+ and TH+ neurons was characterized by a higher number of methylated SST+ neurons and a lower number of methylated TH+ neurons. To investigate whether different photoperiods were correlated with altered overall levels of DNMT3a in the PaVN, we quantified the total number of DNMT3a+ cells per 200 x 200 µM ROI. Since animals at 1 and 3 months were the only age groups to exhibit significant changes, DNMT3a quantification was divided into two groups; younger (1 and 3 months) and older (12 and 18 months). Our quantification showed no changes in the total number of DNMT3a+ cells in the PaVN across photoperiods when compared within age groups (Figure 4F,G), suggesting simultaneous and opposing up- and down-regulation of DNA methylation in different subpopulations of neurons.

**Figure 4.**
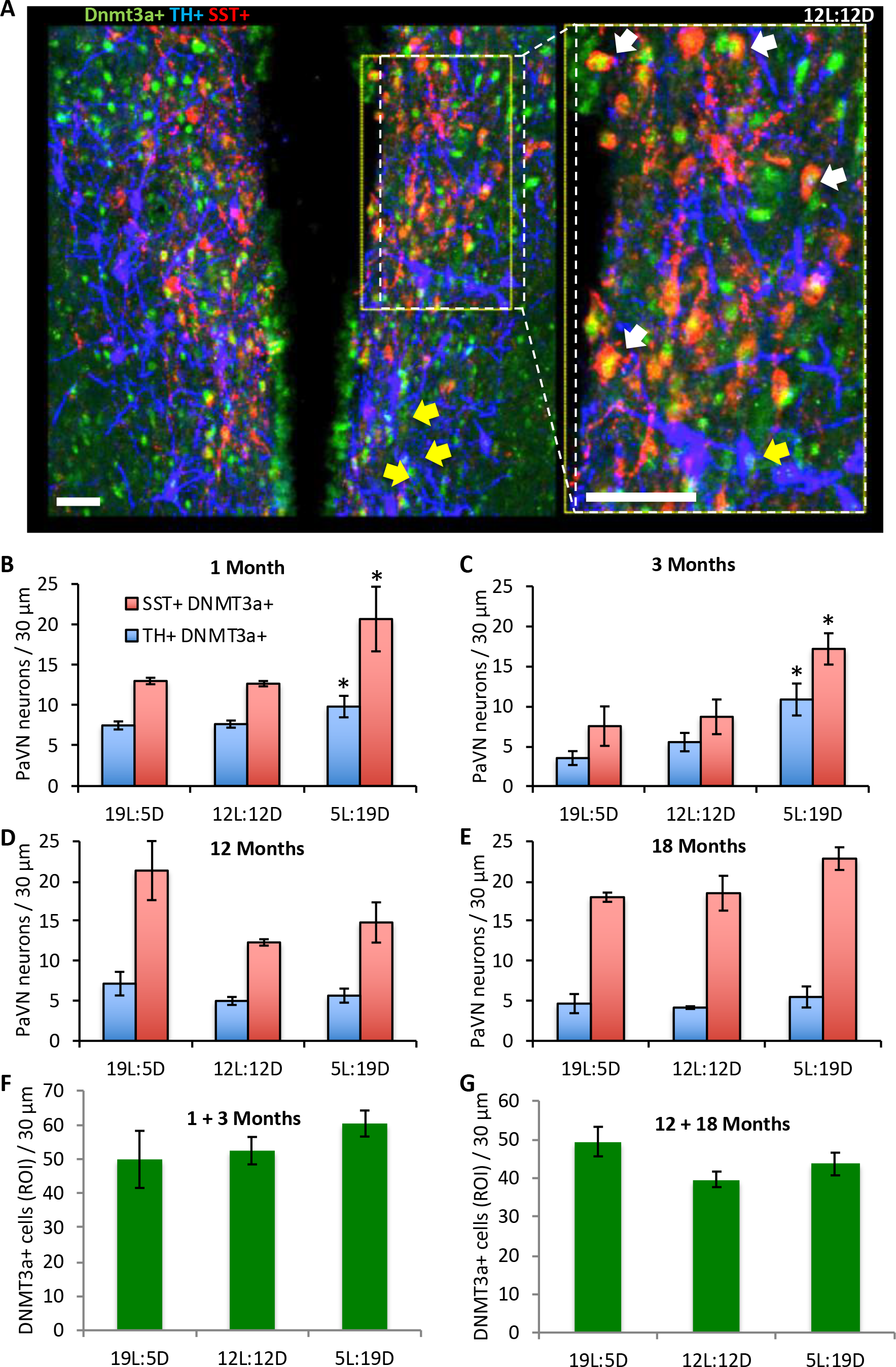
DNA methylation is age-dependent and increases with exposure to short-day photoperiod. (**A**) Immunofluorescence micrograph showing a representative image of the PaVN (control photoperiod) immunostained to label TH+ (blue), SST+ (red), and Dnmt3a+ (green) neurons. Scale bar, 100 µm. The inset (*right*) shows a higher magnification of the same region of the PaVN (*left*, dashed line) and SST+ Dnmt3a+ (white arrows) and TH+ Dnmt3a+ (yellow arrows) neurons. (**B,C**) An increase in DNA methylation in both TH+ and SST+ neurons following short-day photoperiod exposure was observed in animals at 1 & 3 months (*, p=≤ 0.01) of age (n=3/group). (**D,E**) The correlation of Dnmt3a level with photoperiod was lost in 12 and 18 month old rats. **(F,G)** There were no overall changes in the total number of DNMT3a+ neurons (ROI) across photoperiods when compared within age-matched groups (N = 3 animals / group).

### Reciprocal H3 acetylation in SST+/TH+ cells following short-day photoperiod

To investigate the epigenetic mechanisms of DNA accessibility associated with photoperiod-induced neurotransmitter switching, we performed immunohistochemistry to label PaVN TH+ and SST+ neurons with AcetylH3, a marker for histone acetylation at the N-terminus (Figure 5A-C). To this aim we focused on 3-month old rats, since they display high photoperiod-dependent plasticity as well as Dnmt3a-modification in both SST+ and TH+ phenotypes. Our data showed reciprocal H3 acetylation in TH+ versus SST+ neurons of the PaVN following short-day exposure (Figure 5C, 5D) (TH, t_(8)_ = 3.41, p = 0.0092, unpaired; SST, t_(8)_ = 3.57, p = 0.0073, unpaired). Specifically, short-day exposure increased the number of TH+ cells and decreased the number of SST+ cells acetylated at histone 3, relative to balanced-day controls (Figure 5B, 5D). As observed with the methylation data, no difference was found in the total number of SST+ and TH+ neurons exhibiting histone 3 acetylation after long-day photoperiod exposure, compared to balanced-day controls (Figure 5A, 5D). To investigate whether overall levels of H3 acetylation in the PaVN were correlated with different photoperiods, we quantified the total number of AcetylH3+ cells per 200 x 200 µM ROI. Quantification showed no changes in the total number of AcetylH3+ cells in the hypothalamic PaVN across photoperiods (Figure 5E), suggesting differential up- and down-regulation in histone acetylation in selected cell clusters or networks. Such circuit-specific epigenetic regulation could be detected only when the analysis included cellular specificity.

**Figure 5:**
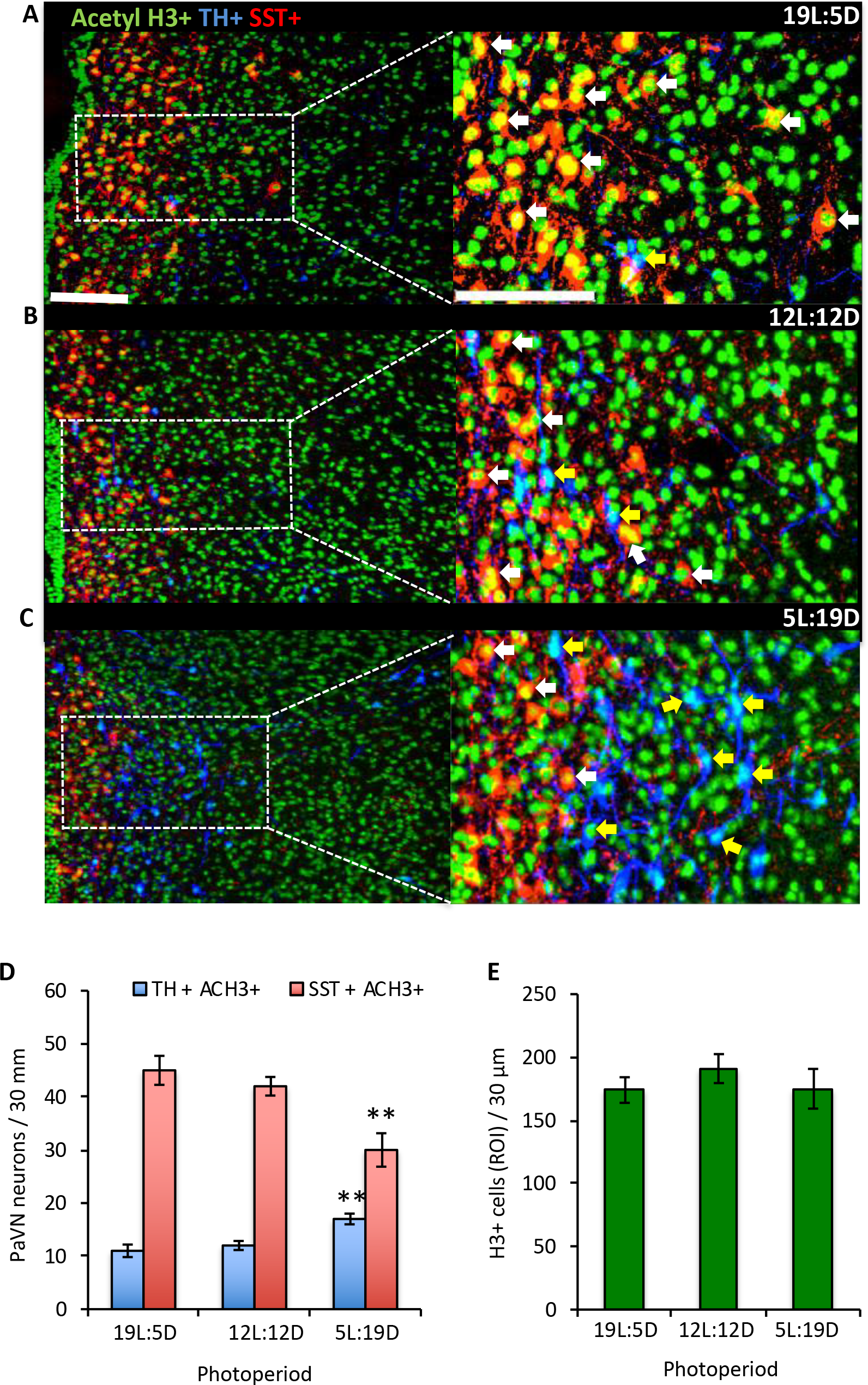
Short-day photoperiod induces reciprocal levels of histone-3 acetylation in SST+ and TH+ neurons. (**A-C**) Confocal micrographs showing representative images of the PaVN of 3-month old rats exposed to long-, balanced-, and short-day photoperiods. Tissue was immunostained to label TH+ (blue), SST+ (red), and Acetyl H3+ (green) neurons. Scale bar, 100 µM. The insets (*right*) show a higher magnification of the same regions of the PaVN (*left*, dashed line) with SST+ Acetyl H3+ (white arrows) and TH+ Acetyl H3+ (yellow arrows) neurons. (**D**) While the number of H3-acetylated SST+ and TH+ neurons increased following short-day photoperiod exposure, there was no significant change after exposure to a long-day photoperiod when compared to controls (n=5 animals / group). (**E**) There was no overall change in the total number of histone-3 acetylated neurons (ROI) across photoperiods (N = 5 animals / group). Scale bars, 100 µm. **, p=≤ 0.001.

## DISCUSSION

This study extends our previous findings on photoperiod-induced neurotransmitter respecification in the adult brain (Dulcis et al., 2013) by determining how this novel form of neuroplasticity is maintained or extinguished with aging. It also investigates possible post-translational epigenetic mechanisms underlying this phenotypic switch in neurotransmitter expression. We found clear age-dependent effects on photoperiod-induced neurotransmitter respecification as well as distinct changes in DNA methylation and histone acetylation in neurons identified by their neurotransmitter expression.

Our results reveal that age plays an important role in the ability of the nervous system to adapt to changes in photoperiod by displaying neurotransmitter plasticity. We found that young rats showed the most robust response to altered photoperiods by up or down-regulating the number of TH+ neurons in the PaVN. However, at 12 months of age, dopaminergic plasticity is displayed only in response to long-day photoperiod exposure (a stressor for nocturnal animals); the ability to increase the numbers of TH+ neurons when exposed to short-day photoperiod has been lost. Since increased levels of dopamine have been shown to mediate anxiolytic and anti-depressive effects when animals were tested for stress responses (Dulcis et al., 2013), our findings indicate that aging animals are less responsive to positive stimuli while still susceptible to stressors. This result is further supported by the finding that older rats (12-15 months) are more vulnerable to stress and are more susceptible to developing anhedonia than younger rats (3-5 months) (Herrera et al., 2008). The increased prevalence of anhedonia among older rats was associated with neuroendocrine changes that occur with aging. Interestingly, we found that TH and SST plasticity did not follow the same time course of decline; SST switching is retained in both photoperiod conditions throughout 12 months of age. This result indicates that the factors mediating age-dependent gene regulation in response to seasonal changes in photoperiod affect neurotransmitter expression differently. At 18 months, both TH and SST plasticity were abolished. The decline of transmitter plasticity in the aging brain may be linked to the documented age-related decrease in the incidence rates of SAD (Kurlansik and Ibay, 2012). The reciprocal shift in TH and SST expression identifies a mechanism by which neural networks can adapt their functions in response to stressful or pleasant environments (Dulcis et al., 2013; Vogels et al., 2011) and may lead to non-pharmaceutical treatments for age-related pathologies associated with dopamine dysfunction.

Other forms of synaptic plasticity such as LTP and LTD are also impacted during aging (Kumar and Foster, 2018; Norris et al., 1996). Much research has focused on the hippocampus, a brain region critically involved in learning and memory, which is particularly susceptible to dysfunction during senescence. Brain aging is associated with altered glutamatergic neurotransmission and Ca^2+^ dysregulation (Kumar and Foster, 2018) that are linked to cognitive impairment. Aged animals show enhanced induction of LTD at hippocampal CA3-CA1 synapses, as a result of differences in calcium homeostasis between young and old rats (Norris et al., 1996). Deficits in synaptic plasticity during normal aging are also attributable to defects in AMPAR trafficking (Norris et al., 1996). In addition, alterations in the size and stability of spines and boutons have been observed in the cortex during brain aging (Mostany et al., 2013). These structural changes result in weaker synapses that are less capable of short-term plasticity in aged individuals reducing circuit function adaptability to a changing environment. The broad umbrella of mechanisms through which aging can negatively influence cognition now includes the impact of age-dependent decline and loss of neurotransmitter switching in addition to reduction in synapse strength and number.

The CNS is constantly being shaped by neural activity throughout the lifetime and epigenetic modifications are important regulators of neural plasticity (Tognini et al., 2015). To examine whether epigenetic modification in TH+ and SST+ neurons involved in photoperiod-induced transmitter switching changed with aging, we investigated the patterns of DNA methylation and histone acetylation in hypothalamic PaVN neurons.

Because DNMT1 methylates hemimethylated DNA to maintain existing methylation patterns (Booth and Brunet, 2016; Jones and Liang, 2009) while DNMT3a methyltransferase introduces *de novo* methylation into otherwise unmodified DNA (Moore et al., 2013), we investigated the expression of DNMT3a as a methylation marker in response to altered photoperiods. We found that younger rats, at 1 and 3 months of age, had an increase in DNA methylation of TH+ and SST+ neurons in the PaVN following the short-day photoperiod exposure only. Intriguingly, *de novo* DNA methylation patterns in this photoperiod revealed an increase in both TH+ and SST+ neurons instead of differential expression patterns predicted from their reciprocal induction of transmitter switching. DNMT3a activity at non-promoter regions causes increased expression of neurogenic genes by hindering cytosine-phosphate-guanine (CpG) binding and gene repression (Lardenoije et al., 2015). While methyltransferases are typically associated with transcriptional silencing, they can also activate neural plasticity (Laplant et al., 2010; Maze et al., 2010). Given that methylation has site-specific opposing effects on gene expression, further work needs to be done to determine the mechanism through which DNMT3a acts in TH- and SST-expressing neurons. As age increased, the pattern of photoperiod-dependent DNMT3a localization in both phenotypes was lost and instead marked by an overall increase in *de novo* methylation in SST+ cells across all photoperiods. Methylation patterns are known to change with aging and can result in local promoter-specific hypermethylation (Johnson et al., 2012; Jung and Pfeifer, 2015). The overall increase in DNMT3a expression in SST+ cells of the PaVN may be attributable to polycomb loss since unmethylated regions bound by polycomb complexes have been shown to become methylated with aging (Jung and Pfeifer, 2015; Cedar and Bergman, 2012). This process would increase access to DNA and allow methylation levels to increase over time, resulting in a loss of plasticity. However, in younger rats, DNMT3a may be inhibiting CpG binding (Lardenoije et al., 2015) to increase plasticity and increase neurogenic gene activity. Total DNMT3a expression was quantified in the PaVN and compared across photoperiods within age groups and revealed no differences in overall levels, pointing to changes at the circuit level rather than overall expression.

As with changes in DNMT3a expression in younger rats, significant changes in histone 3 (H3) acetylation were observed only following short-day photoperiod exposure. However, TH+ and SST+ neurons exhibited a reciprocal pattern of H3 acetylation different from that of *de novo* methylation. Short-day photoperiod exposure increased H3 acetylation in TH+ neurons and decreased H3 acetylation in SST+ cells. Because acetylation is typically associated with gene transcription, the findings were consistent with our hypothesis that acetylation levels of H3 would match the patterns of dopamine and SST neurotransmitter switching following short-day exposure. Histones H3 and H4 are particularly important in gene expression, because posttranslational modifications happening at these two histones are associated with transcriptional states (Annunziato, 2002; Yan and Boyd, 2006). For instance, recruitment of Nurr1, a transcription factor crucial for midbrain dopaminergic neuron development, to the promoter of TH gene has been shown to be enhanced by depolarization along with an increase of histone 3 acetylation in the Nurr1 binding regions of TH promoter (He et al., 2011). Total H3 acetylation quantified in the hypothalamus and compared across photoperiods revealed no differences in overall levels. However, identifying a reciprocal histone acetylation pattern and an increase in de novo methylation in TH+ and SST+ neurons, in spite of a constant level of total acetylation and methylation, underscores the importance of studying epigenetic mechanisms at the circuit level and quantifying identified cell types. Changes in SST in the PaVN are matched by corresponding changes in mRNA levels and involve translation of *de novo* SST mRNA (Dulcis et al., 2013; Arancibia et al., 2000). While neurons are losing SST and being recruited to newly express TH, both translational silencing markers and pro-transcriptional markers may work in concert to stabilize the epigenome of these cells according to photoperiod. In summary, this study provides a map of TH/SST neuroplasticity across aging, shows an age-dependent expression of methylation patterns across photoperiods and demonstrates a reciprocal H3 acetylation in neurons that switch their transmitter in the adult brain. Further work will more fully elucidate the roles of DNMT3a and H3 acetylation in sensory-stimulus induced neurotransmitter respecification via protein-DNA interactions.

## Acknowledgments

We thank Dr. F. Telese for her expert advice on qPCR protocols. This work was supported by W. M. Keck Foundation to N.C.S and D.D. (Grant # 20132777).

